# Automatic vegetation identification in Google Earth images using a convolutional neural network: A case study for Japanese bamboo forests

**DOI:** 10.1101/351643

**Authors:** Shuntaro Watanabe, Kazuaki Sumi, Takeshi Ise

## Abstract

Classifying and mapping vegetation are very important tasks in environmental science and natural resource management. However, these tasks are not easy because conventional methods such as field surveys are highly labor intensive. Automatic identification of target objects from visual data is one of the most promising ways to reduce the costs for vegetation mapping. Although deep learning has become a new solution for image recognition and classification recently, in general, detection of ambiguous objects such as vegetation still is considered difficult. In this paper, we investigated the potential for adapting the chopped picture method, a recently described protocol for deep learning, to detect plant communities in Google Earth images. We selected bamboo forests as the target. We obtained Google Earth images from three regions in Japan. By applying the deep convolutional neural network, the model successfully learned the features of bamboo forests in Google Earth images, and the best trained model correctly detected 97% of the targets. Our results show that identification accuracy strongly depends on the image resolution and the quality of training data. Our results also highlight that deep learning and the chopped picture method can potentially become a powerful tool for high accuracy automated detection and mapping of vegetation.

## INTRODUCTION

Classifying and mapping vegetation are essential tasks for environmental science research and natural resource management^1^. Traditional methods (e.g., field surveys, literature reviews, manual interpretation of aerial photographs), however, are not effective for acquiring vegetation data because they are labor intensive and often economically expensive. The technology of remote sensing offers a practical and economical means to acquire information on vegetation cover, especially over large areas^2^. Because of its systematic observations at various scales, remote sensing technology potentially can enable classification and mapping of vegetation at high temporal resolutions.

Detection of discriminating visual features is one of the most important steps in almost any computer vision problem, including in the field of remote sensing. Since conventional methods such as support vector machines^3^ require hand-designed, time-consuming feature extraction, substantial efforts have been dedicated to development of methods for the automatic extraction of features. Recently, deep learning has become a new solution for image recognition and classification because this new method does not require the manual extraction of features.

Deep learning^4,5^ is one type of machine learning technique that uses algorithms inspired by the structure and function of the brain called artificial neural networks. Deep learning involves the learning of features and classifiers simultaneously, and it uses training data to categorize image content without a priori specification of image features. Among all deep learning-based networks, the convolutional neural network (CNN) is the most popular for learning visual features in computer vision applications including remote sensing. Recent research has shown that CNN is effective for diverse applications^4-7^. Given its success, deep learning has been used intensively in several distinct tasks for different academic and industrial fields including plant science. Recent research has shown that the deep learning technique can successfully detect plant disease, correctly classify the plant specimens in a herbarium ^8-10^

Deep learning is a promising technology also in the field of remote sensing^11,12^. Recently, Guirado et al., (2017)^13^ demonstrated that the deep learning technique successfully detect plant species of conservation concern and it provides better results than the conventional object detection methods. However, application of deep learning to vegetation mapping are not sufficient yet because vegetation in the aerial image often shows ambiguous and amorphous shape, and automatic object identification including deep learning tends not to work well on such objects.

Recently, Ise et al., (2018)^14^ developed a method to extract relevant characteristics from ambiguous and amorphous objects. This method dissects the images into numerous small squares and efficiently produces the training images. By using this method, Ise et al. (2018)^14^ correctly classified three moss species and “non-moss” objects in test images with an accuracy of more than 90%.

In this paper, we investigated the potential for adapting a deep learning model and the chopped picture method to automatic vegetation detection in Google Earth images, and bamboo forests were used as the target. In recent years, bamboo has become invasive in Japan. The bamboo species moso (*Phyllostachys edulis*) and madake (*P. bambusoides* Siebold) are the two main types of exotic bamboo. Since the 1970s, the bamboo industry in Japan had declined as a result of cheaper bamboo imports and heavy labor costs^15^. Consequently, many bamboo plantations were left unmanaged, which led to the eventual invasion of adjacent native vegetation ^16-18^.

In this study, we specifically addressed the following questions: 1) how does the resolution of images affect the accuracy of detection; 2) how does the chopping size of training images affect the accuracy of detection; and 3) can a model that learned in one geographical location work well for a different location?

## MATERIALS AND METHODS

### Target area and image acquisition

In this study, we chose three regions (Sanyo-Onoda, Ide, and Isumi) in Japan to conduct the analyses (Figure 1). We used Google Earth as the source of imagery. From a given sampling location, we obtained the images at zoom levels of 1/500 (~0.13 m/pixel spatial resolution), 1/1000 (~0.26 m/pixel spatial resolution), and 1/2500 (~0.65 m/pixel spatial resolution).

**FIGURE 1.**
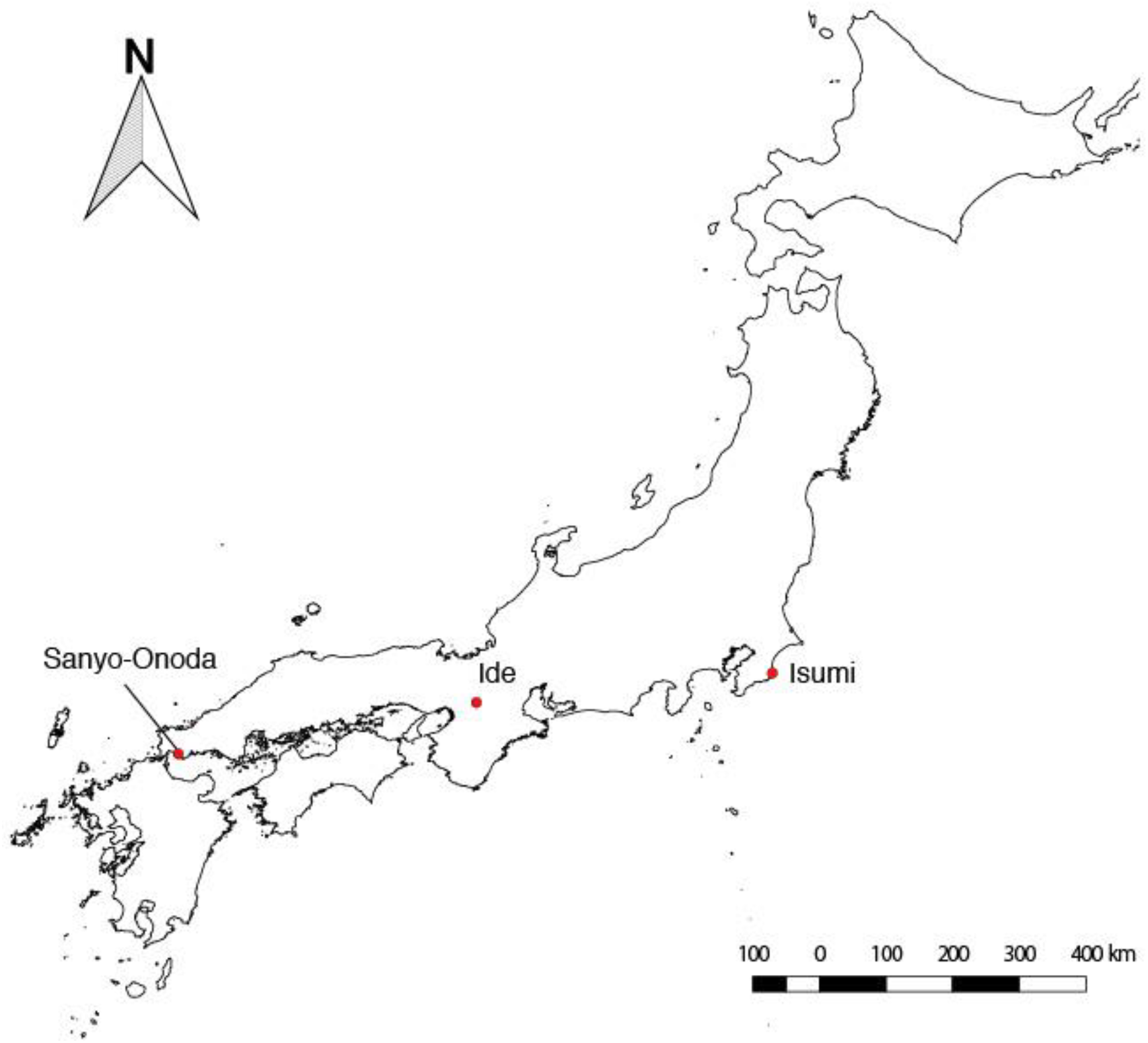
Target regions of this research.

### Methods and background concepts for the neural networks

In this study, we employed convolutional neural networks (CNN; Figure 2). A CNN is a special type of feedforward neural network that consists of a convolutional layer and pooling layer.

**FIGURE 2.**
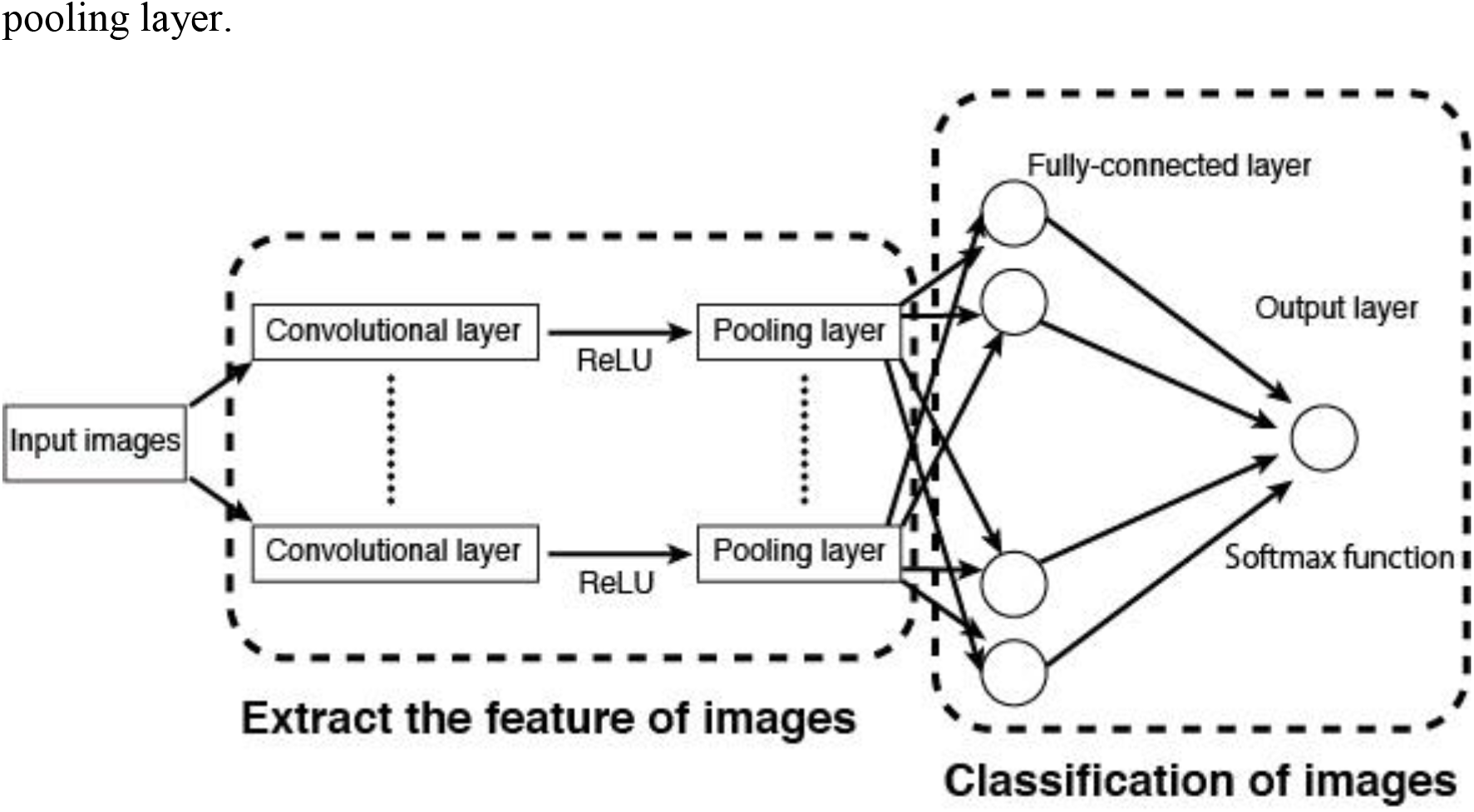
Schematic diagram of the convolutional neural networks.

A feedforward neural network is an artificial neural network wherein connections between the nodes do not form a cycle. These networks, which conduct modeling similar to the neuron activity in the brain, are generally presented as systems of interconnected processing units (artificial neurons) that can compute values from inputs leading to an output that may be used on further units. Artificial neurons are basically processing units that compute some operation over several input variables and, usually, have one output calculated through the activation function. Typically, an artificial neuron has a weight *w_i_* that represents the degree of connection between artificial neurons, some input variables *x_i_*, and a threshold vector *b*. Mathematically, the total input and output of artificial neurons can be described as follows:

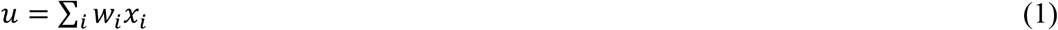

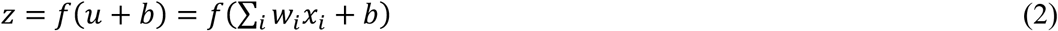
where *u*, *z*, *x*, *w*, and *b* represent the total input, output, input variables, weights, and bias, respectively. *f*(∙) denotes an activation function; a nonlinear function such as a sigmoid, hyperbolic, or rectified linear function is provided in *f*(∙). We employed a rectified linear function as the activation function, and this function is referred to as the Rectified Linear Unit (ReLU). The definition of ReLU is shown in the following equation:

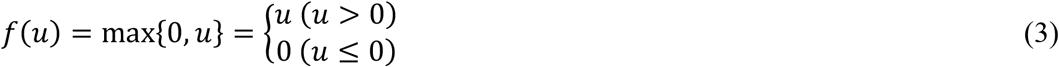

As mentioned above, a CNN is a special type of feedforward neural network that is usually used in image classification and identification. A CNN consists of a convolutional layer and pooling layer. The convolutional layer plays a role in capturing the features from the images. In this process, a fixed-sized window runs over the image and extracts the patterns of shades of colors in the image. After each convolutional layer, there are pooling layers that are created in order to reduce the variance of features, which is accomplished by computing some operation of a particular feature over a region of the image.

The pooling layer has the function of reducing the position sensitivity of the feature that is extracted at the convolution layer so that the output amount of the pooling layer does not change even when the position of the feature amount extracted by the convolution layer is shifted within the image. Two operations may be realized on the pooling layers, namely, max or average operations, in which the maximum or mean value is selected over the feature region, respectively. This process ensures that the same results can be obtained, even when image features have small translations or rotations, and this is very important for object classification and detection. Thus, the pooling layer is responsible for sampling the output of the convolutional one and preserving the spatial location of the image, as well as selecting the most useful features for the next layers.

After several convolutional and pooling layers, there are fully connected ones, which take all neurons in the previous layer and connect them to every single neuron in its layer.

Finally, following all of the convolution, pooling, and fully connected layers, a classifier layer may be used to calculate the class probability of each instance. We employed the softmax function in this layer. The softmax function calculates the probabilities of each target class over all possible target classes. The softmax function is written as follows:

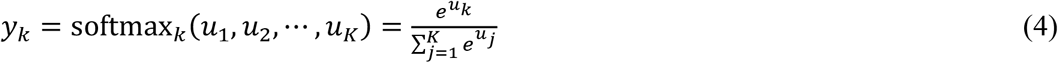
where *k* represents the number of the output unit and *u* represents input variables.

In order to evaluate the performance of the network, a loss function needs to be defined. The loss function evaluates how well the network models the training dataset. The goal of the training is to minimize the error of the loss function. Eq. (5) presents the cross entropy of the softmax function that was employed in this study:

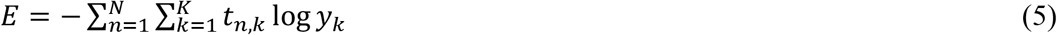
where *t* is the vector for the training data, *K* represents the possible class, and *N* represents the total number of instances.

### Approach

A schematic diagram of our approach is shown in Figure 3. We prepared the training data by using the chopped picture method ^14^. First, in this method, we collected the images that were (1) nearly 100% covered by bamboo and (2) not covered by bamboo. Next, we “chopped” this picture into small squares with 50% overlap both vertically and horizontally.

**FIGURE 3.**
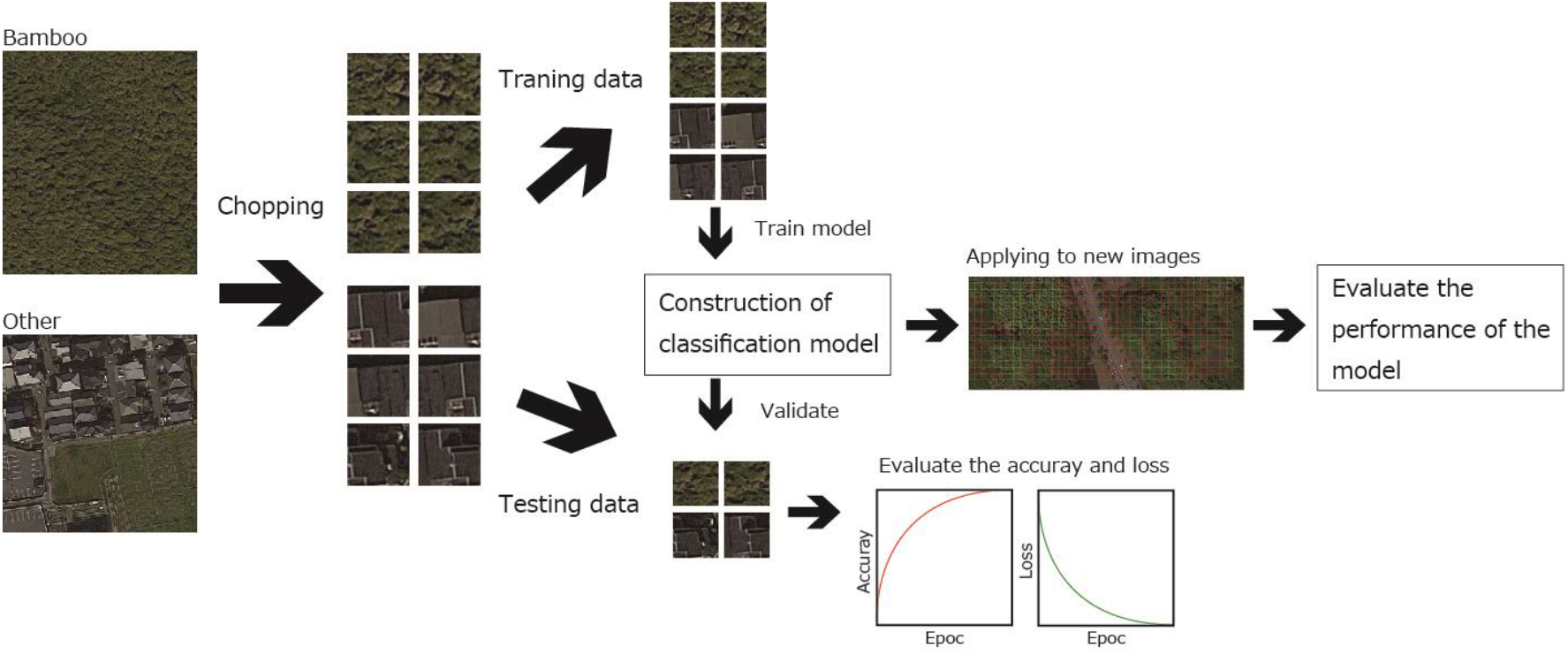
Schematic diagram of the research approach. This figure was generated using data from Google Earth image (Image data: ©2018 CNES/Airbus & Digital Globe).

We made a model for image classification from a deep CNN for the bamboo forest detection. As opposed to traditional approaches of training classifiers with hand-designed feature extraction, the CNN learns the feature hierarchy from pixels to classifiers and trains layers jointly. We used the final layer of the CNN model for detecting the bamboo coverage from Google Earth images. To make a model for object identification, we used the deep learning framework of NVIDIA DIGITS^19^. We used 75% of the obtained images as training data and the remaining 25% as validation data. We used the LeNet network model^20^. The model parameters implemented in this study included the number of training epochs (30), learning rate (0.01), train batch size (64), and validation batch size (32).

### Evaluation of the learning accuracy

Validation of the model in each learning epoch was conducted by using the accuracy and loss function obtained from cross validation images. The accuracy indicates how accurately the model can classify the validation images. Loss represents the inaccuracy of the prediction of the model. If model learning is successful, loss (val) is low and accuracy is high. However, when loss (val) becomes high during learning, this indicates that over fitting is occurring.

### Evaluation of the model performance

We obtained 10 new images, which were uniformly covered by bamboo forest or objects other than bamboo forest, from each study site. Next, we re-sized the images by using the chopped picture method. Third, we randomly sampled 500 images from the re-sized images. Finally, we applied the model to the sampled images and evaluated the classification accuracy. To evaluate the performance of the model, we classified the classification results into the following four categories: true positive (*TP*), false positive (*FP*), false negative (*FN*), and true negative (*TN*). Next, we calculated the classification accuracy, recall rate, and precision rate by using the following equations:

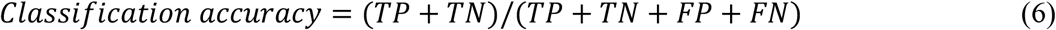

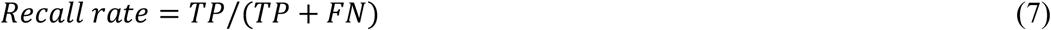

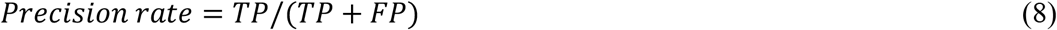

### Testing the effects of image resolution on the classification accuracy

To quantify the effects of image resolution on the accuracy of detection, we obtained images at a zoom level of 1/500 (~0.13 m/pixel spatial resolution), 1/1000 (~0.26 m/pixel spatial resolution), and 1/2500 (~0.65 m/pixel spatial resolution) from each study site. Next, we applied the chopped picture method. To adjust the spatial extent of each chopped image, we chopped 56 pixels for the 1/500 level, 28 pixels for the 1/1000 level, and 14 pixels for the 1/2500 level. After constructing the model, we applied the model to the new images and calculated the classification accuracy, recall rate, and precision rate.

### Testing the effects of chopping grid size on the classification accuracy

To quantify the effects of the spatial extent of the chopping grid on the accuracy of detection, we chopped the 1/500 images at each study site for three types of pixel sizes (84, 56, 28). After constructing the model, we applied the model to new images and calculated the classification accuracy, recall rate, and precision rate.

### Transferability test

Given the large amount of variation in the visual appearance of bamboo forest across different cities, it is of interest to study to what extent a model learned on one geographical location can be applied to a different geographical location. As such, we performed experiments in which we trained a model for one (or more) cities, and then, we applied the model to a different set of cities. Performance of the model was evaluated by the classification accuracy, recall rate, and precision rate.

## RESULTS

### Fluctuation of accuracy and loss during the learning epochs

The accuracy for classifying the validation data of the final layer ranged from 94% to 99%. Loss values for the validation data ranged from 0.008 to 0.214 (Figure 4). Values of accuracy increased and loss decreased following the learning epochs (Figure 4). These results suggest the all of the models were not overfit to the datasets and successfully learned the features of chopped pictures.

**FIGURE 4.**
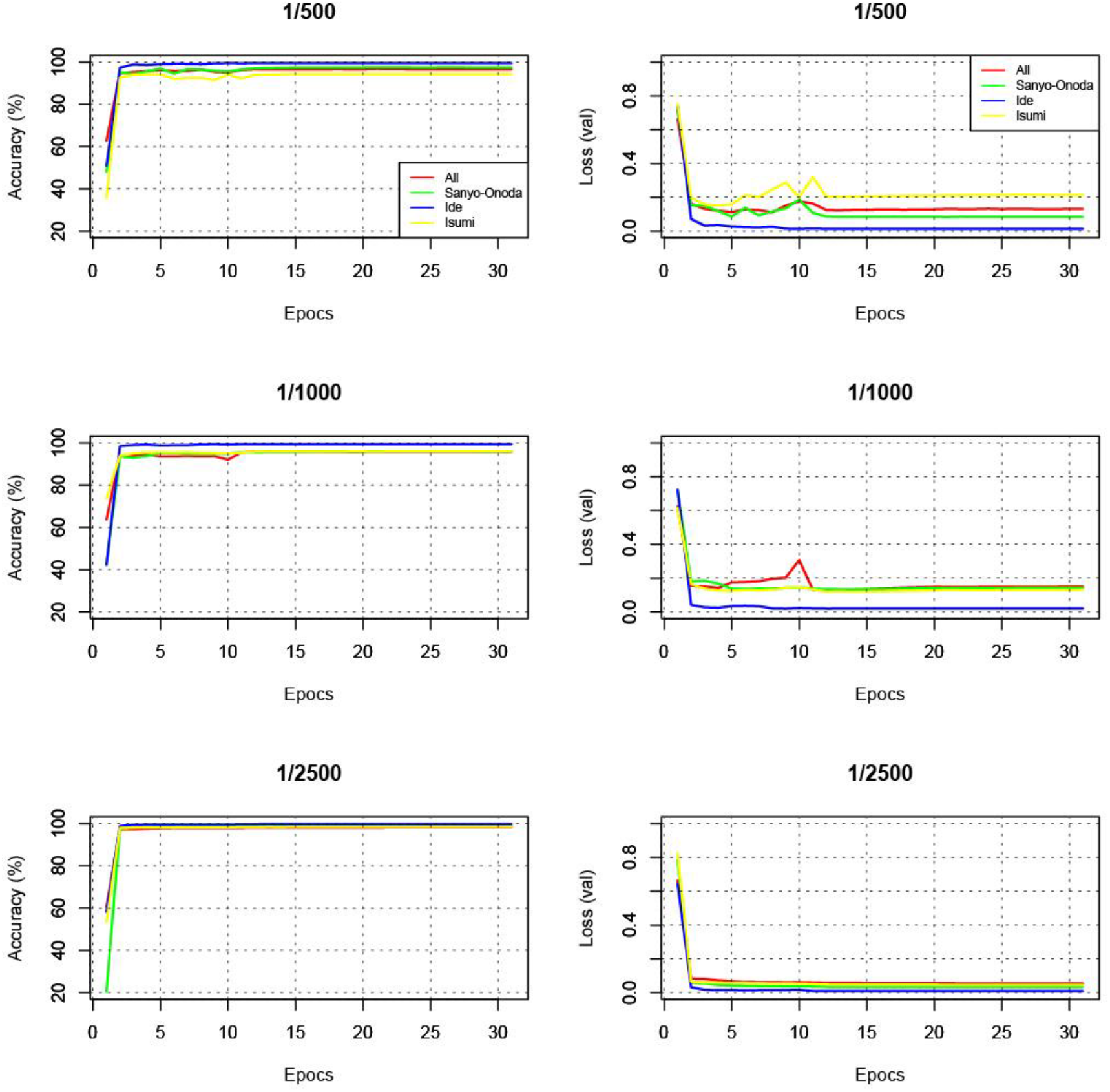
Accuracy and loss (val) of the model at each learning epoch.

### Effects of image resolution on the classification accuracy

The classification accuracy ranged from 76% to 97% (Figure 5a). The recall rate and precision rate for bamboo forest ranged 52% to 96% and 91% to 99%, respectively (Figure 5b, d). The recall rate and precision rate for objects other than bamboo forest ranged from 92% to 99% and 67% to 96%, respectively (Figure 5c, e). The recall rate for bamboo forest declined following the decline in the image resolution, and it declined dramatically when we used the 1/2500 level (~0.65 m/pixel spatial resolution) (Figure 5a).

**FIGURE 5.**
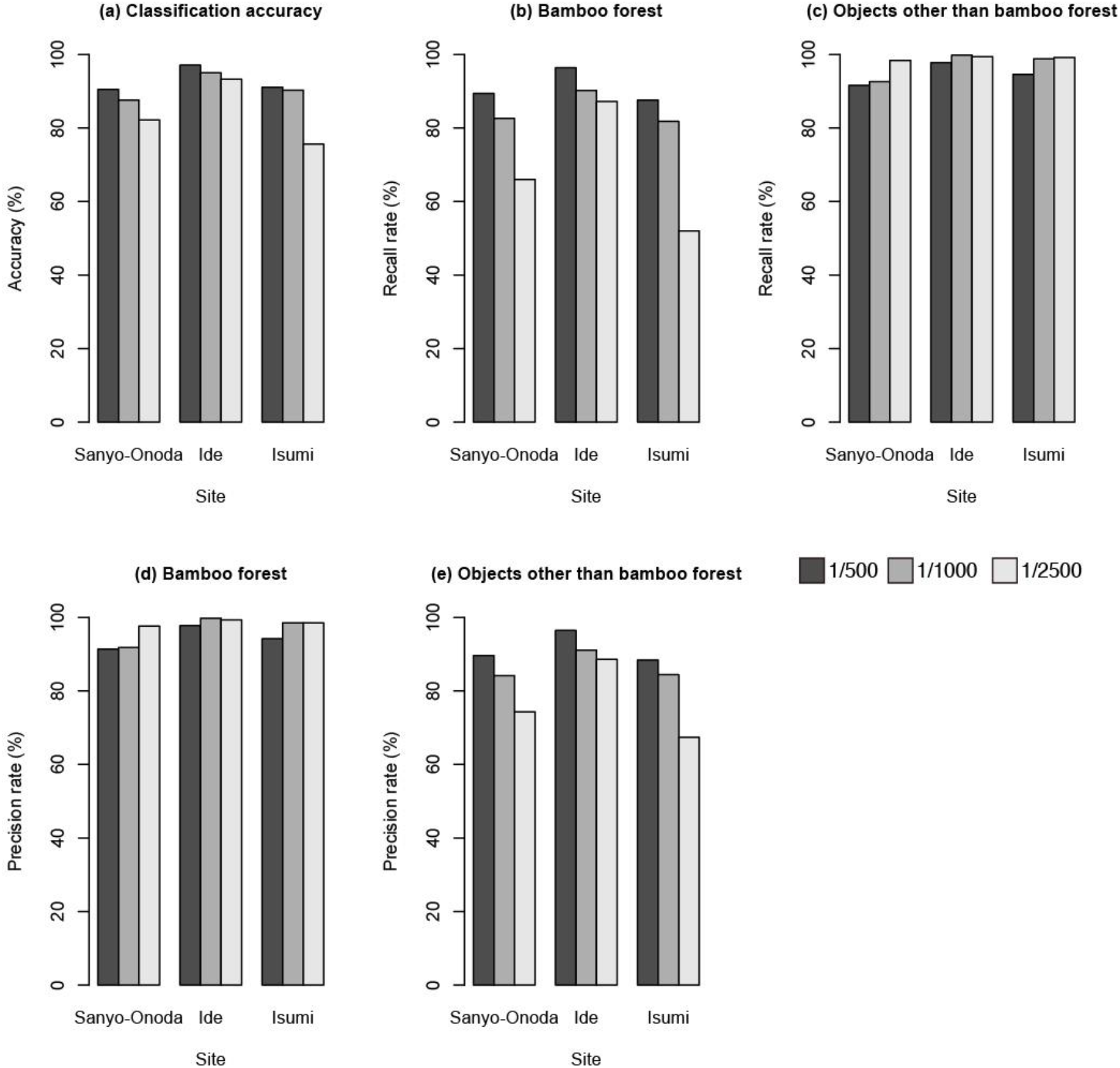
Sensitivity of the image scale versus the test accuracy.

### Effects of chopping grid size on the classification accuracy

The classification accuracy ranged from 85% to 96% (Figure 6a). The recall rate and precision rate for bamboo forest ranged from 79% to 99% and 89% to 98%, respectively (Figure 6b, d). The recall rate and precision rate for objects other than bamboo forest ranged from 88% to 98% and 79% to 99%, respectively (Figure 6c, e). The intermediate-sized images (56 pixel) showed the highest classification accuracy at all sites (Figure 6a). An example of a classification image is shown in Figure 7.

**FIGURE 6.**
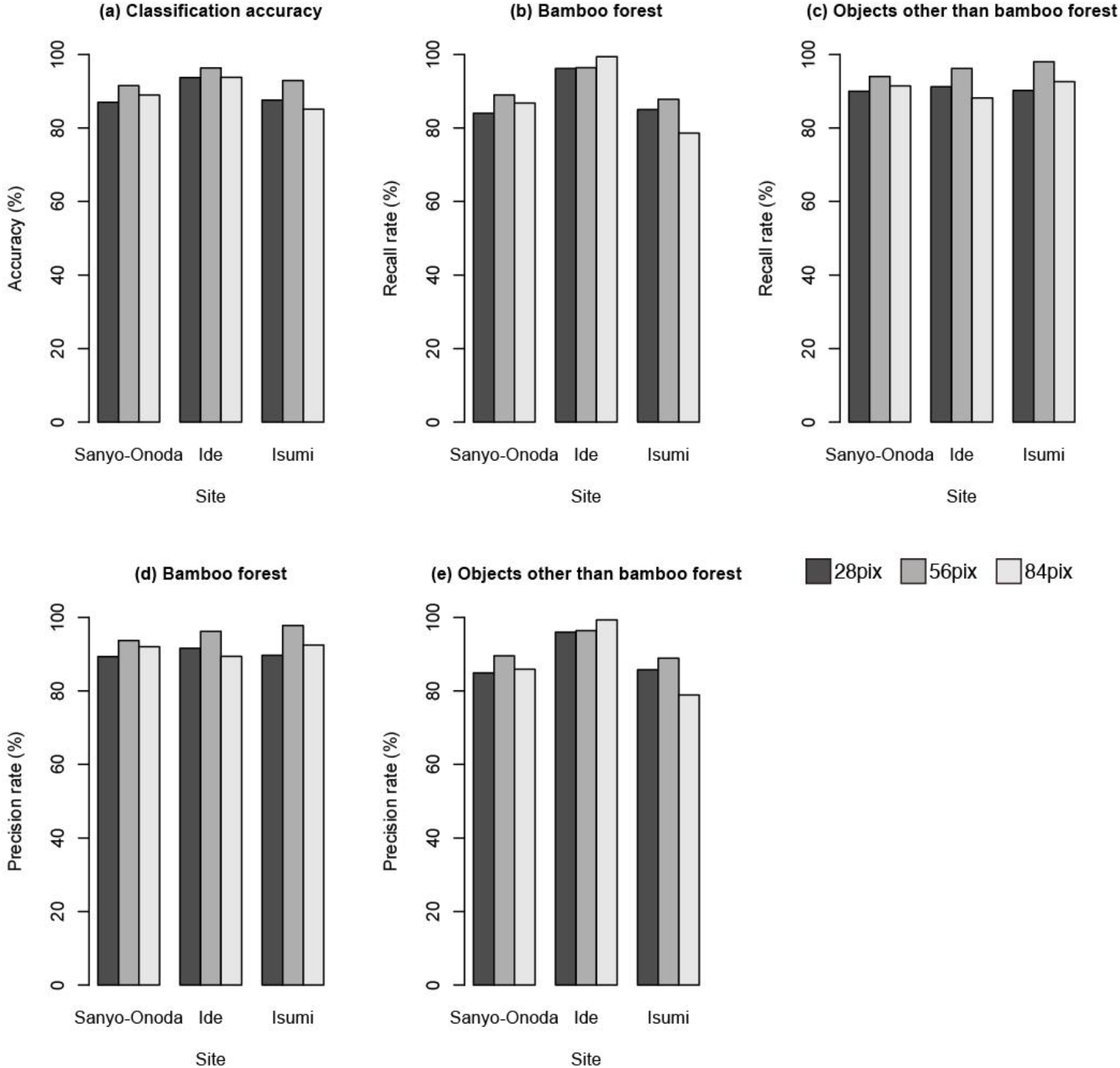
Sensitivity of the pixel size versus the test accuracy.

**FIGURE 7.**
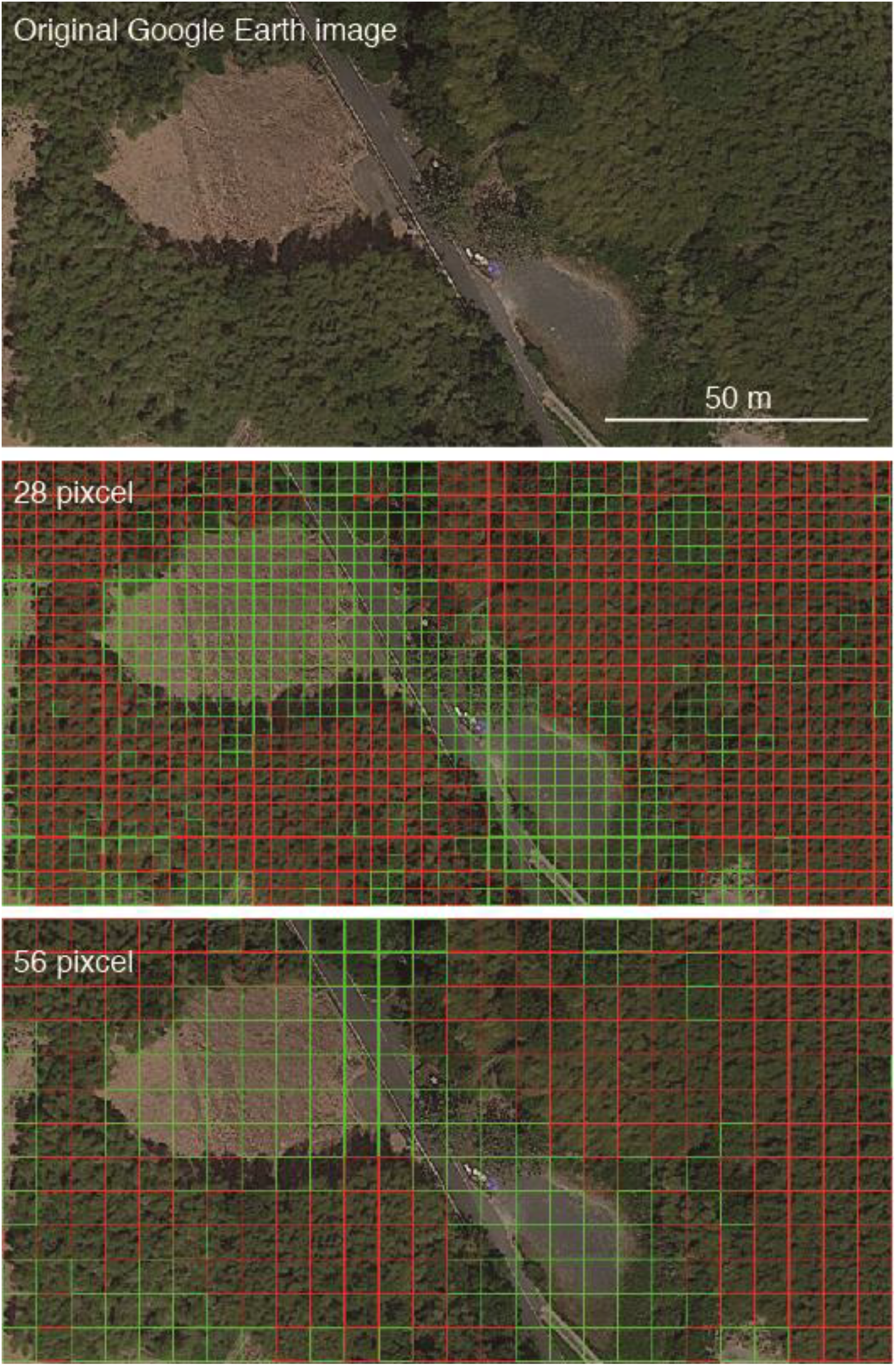
Example of a classification image. Bamboo forests are highlighted by red, and objects other than bamboo are highlighted by green. This figure was generated using data from Google Earth image (Image data: ©2018 CNES/Airbus & Digital Globe).

### Transferability and classification performance

In general, performance was poor when training on samples from a given city and testing on samples from a different city (Figure 8a). When the model that was trained by the images of Isumi city was applied to the other cities, the recall rate was the worst (Figure 8b). Conversely, the model that was trained by the images of Sanyo city showed the highest recall rate (Figure 8b). We noticed that a more diverse set (all) did not yield a better performance when applied at different locations than the models trained on individual cities (Figure 8).

**FIGURE 8.**
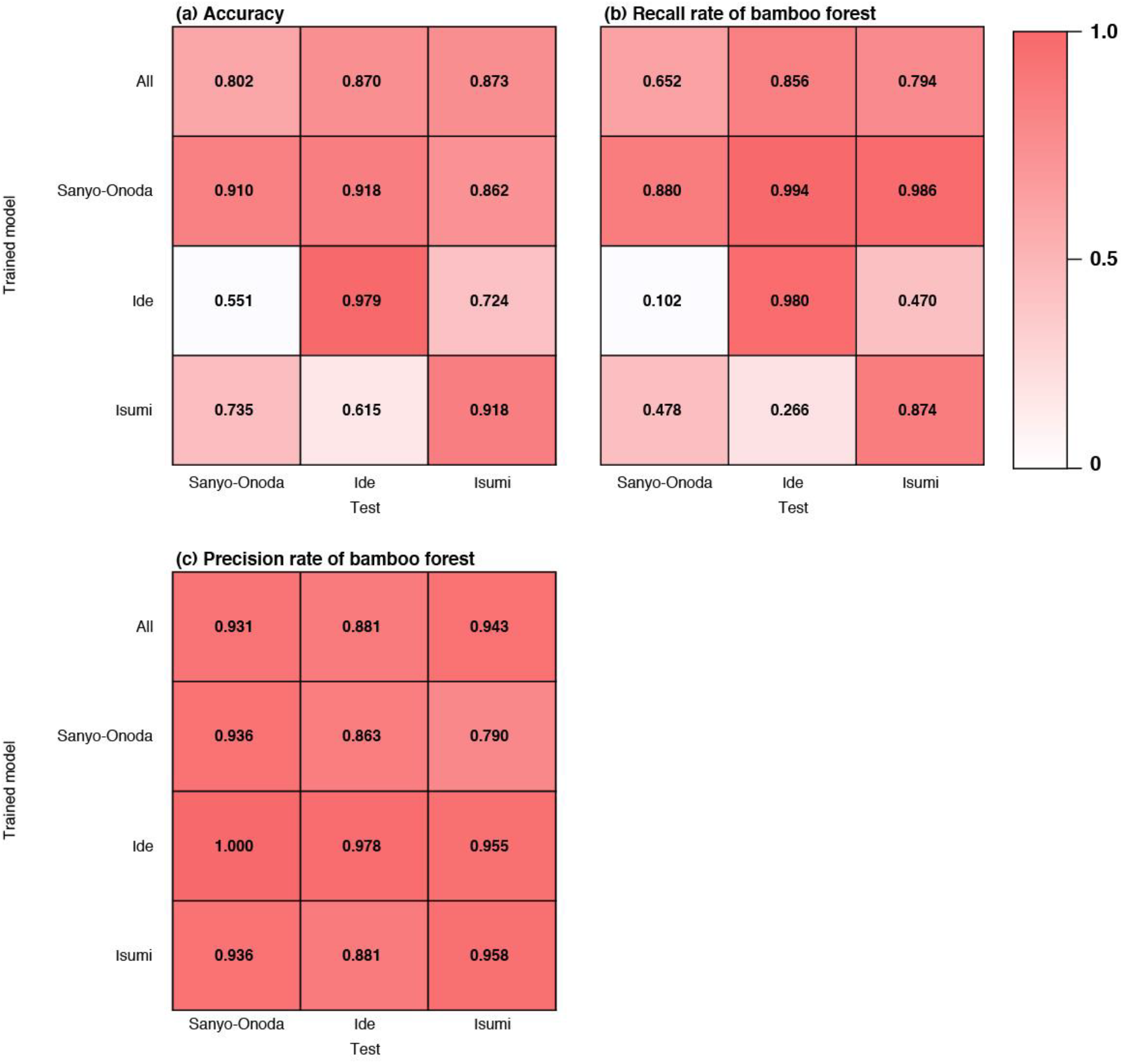
Transferability of the models learned at one location and applied at another.

## DISCUSSION

In this paper, we demonstrated that the chopped picture method and CNN could accurately detect bamboo forest in Google Earth imagery (see Figure 7). Recent research has shown that the deep learning technique is applicable to plant science research ^8-11, 21,22^; however, applications of DA in plant science have been mainly restricted to photographs taken indoors, and applications to plants in the aerial photographs are still limited^11^. To the best of our knowledge, this is the first study to successfully identify plant communities automatically from Google Earth imagery.

Classifying vegetation from remote sensing images generally suffers from several problems, e.g., it can be difficult to establish a specific pattern for each species, given the high intraclass variance, and also to distinguish between different species, given the interclass similarity of distinct species^12,23^. So far, it is generally been difficult to detect ambiguous objects such as vegetation, but our results showed good performance during the detection of bamboo forest from Google Earth images by using the chopped picture method even though we employed the most classical CNN (LeNet). Our results highlight that the chopped picture method and CNN would be a powerful method for high accuracy automated bamboo forest detection and vegetation mapping (see Figure 9).

**FIGURE 9.**
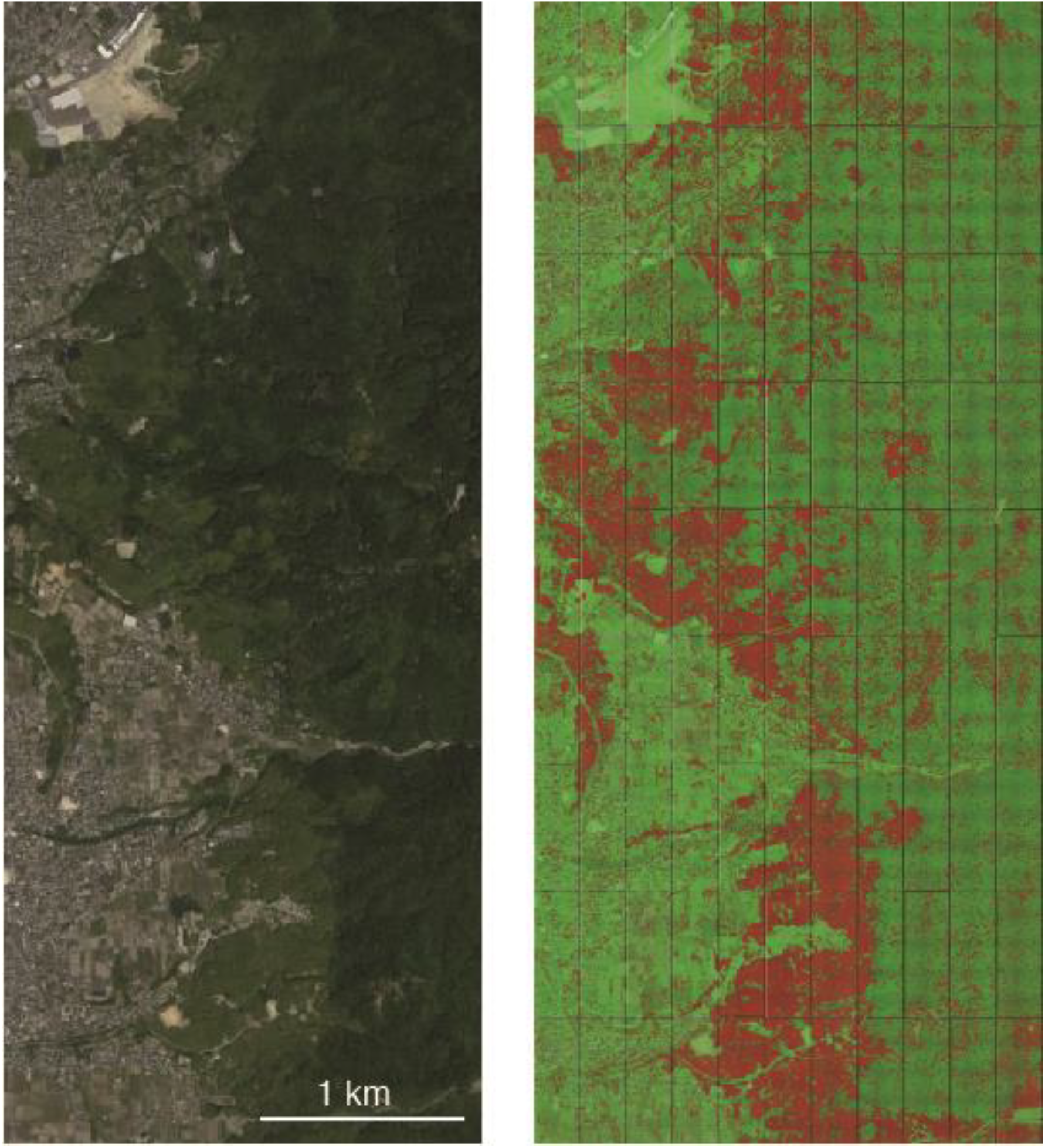
Example of applying the model to the wide area of Ide city. The left image is the original Google Earth image, and the right image shows the results of bamboo forest detection. Bamboo forests are highlighted by red, and objects other than bamboo are highlighted by green. This figure was generated using data from Google Earth image (Image data: ©2018 CNES/Airbus & Digital Globe).

### Effects of image resolution on the classification accuracy

Our results indicate that the image resolution strongly affects the identification accuracy (Figure 4). As the resolution rate decreased, performance of the model also declined (Figure 4).

Especially in the 1/2500 imagery, the recall rate for bamboo forest of Sanyo-Onoda and Isumi city declined to 53% and 64%, respectively (Figure 4b). In contrast, the precision rate for bamboo forest increased as the resolution decreased (Figure 4d). This result means that as the resolution decreases, the model overlooks many bamboo forests; thus, when the image resolution is low, it is difficult to learn the features of the object. This result also suggests that in the deep learning model, misidentification due to false negatives is more likely to occur than misidentification due to false positive as the image resolution declines.

### Effects of chopping grid size on the classification accuracy

Our results indicate that the chopping grid size also affects the performance of the model. The classification accuracy was the highest at the medium pixel size (56 × 56 pixels; Figure 5a). In contrast to the effects of image resolution, the recall rate and precision rate for bamboo forest was also the highest at the medium pixel size except for the recall rate at Ide city (Figure 5b, d).

This result means that if the grid size is inappropriate, both false positives and false negatives will increase.

Increases of the chopping grid size will cause an increase in the number of chopped pictures in which objects other than bamboo and bamboo are mixed. In this paper, because we evaluated the performance of the model by using images that were uniformly covered by bamboo forest or objects other than bamboo forest, the effects of imagery consisting of mixed objects on the classification accuracy could not be evaluated. Evaluation of the classification accuracy for such images will take place in future research.

### Transferability among the models

Results for the transferability tests showed that transferability was generally poor and suggest that the spatial extent of acquisition of training data strongly influences the classification accuracy (Figure 8). The model trained by Sanyo-Onoda city images yielded high recall rates for the images taken at all of the study sites, but the precision rate was lower than that of the other models (Figure 8b, c). This means that the model trained by Sanyo-Onoda city images tends to make false positive mistakes.

Interestingly, transferability was not found to be related to the distance among the study sites (Figure 8). This result indicates that classification accuracy across the model reflects the conditions at the local scale such as the climate at the time when the image was taken. Additionally, even when we applied a model that learned from all training images (all), the performance of the model was not as good as when the training data were obtained within the same city. The same tendencies have been reported in studies that classified land use by using deep learning^24^. This may suggest that increasing the number of training data may also lead to a decrease in the identification accuracy, and it may be difficult to construct an identification model applicable to a broad area.

### Conclusions and future directions

Our results show that the deep learning model presented here can detect bamboo forest from Google Earth images accurately. Our results also suggest that deep learning and the chopped picture method would be a powerful tool for high accuracy automated vegetation mapping and may offer great potential for reducing the effort and costs required for vegetation mapping as well as improving the current status of monitoring the distribution of bamboo. Recently, bamboo expansion has become an important problem in Japan because of its invasiveness^17^. While some research has analyzed the bamboo forest distribution probability on a national scale^25,26^, monitoring of bamboo expansion is still a challenging problem because of labor requirements. Our approach could potentially lead to the creation of a semi or even fully automated system for the monitoring of bamboo expansion.

Our results also suggest that the identification accuracy depends on the image resolution and chopping grid size. Especially, the spatial resolution of training data strongly affects the model performance. Generally, satellite-based remote sensing has been widely studied and applied but suffers from insufficient information due to low resolution images or inaccurate information due to local weather conditions^27^. Our results also show that the performance of the model can be greatly influenced by the spatial extent of the acquired training data and a model learned on one geographical location is difficult to apply to a different geographical location. It remains a future task to develop a model that can be applied over a wide spatial scale.

## ACKNOWLEDGEMENTS

This work was supported by JST PRESTO, Japan (Grant No. JPMJPR15O1).

## Author Contributions

S. W designed the research based on discussion with T. I. S. W and K. S conducted research. S. W and T. I wrote the manuscript.

